# Looks can be deceiving: discordances in phylogeny and morphology within loricate choanoflagellates

**DOI:** 10.1101/2025.07.22.666139

**Authors:** Alex Gàlvez-Morante, Florentine U. Rutaganira, Brian Palenik, Daniel J. Richter

## Abstract

Choanoflagellates are heterotrophic holozoans that are classified into two groups based on their morphology: loricates, which possess a silica-based extracellular structure, and craspedids, which do not. Although the craspedid versus loricate morphological separation is currently supported by their phylogenetic relationship, recent evidence has suggested inconsistencies between morphology and phylogeny within each group. Loricate choanoflagellate taxonomy has historically been based on selected aspects of their lorica morphology, and on their mode of cell division, in which tectiform daughter cells emerge into a lorica synthesized by their mother cell following division, and nudiform daughter cells do not. Here, we characterize two new loricate strains that display unexpected morphological features when compared to their nearest genetic relatives. The strain BEAP0094 very closely matched the 18S ribosomal gene of the tectiform *Pseudostephanoeca paucicostata*, but its morphology clearly differed, due to the absence of the characteristic anterior ring found in all *Stephanoeca* species. Instead, its features resembled more closely those of the *Acanthocorbis* genus, raising the possibility of the existence of either multiple lorica morphologies within the same or very closely related species, or multiple morphological species sharing the same 18S ribosomal gene. The second strain we investigated, BEAP0360, presented a morphological match to *Stephanoeca cauliculata*, but its 18S ribosomal sequence did not, suggesting that different species could share the same lorica architecture. BEAP0360, here described as *Cepoeca plumat*a (n. gen. n. sp.), possesses a key phylogenetic placement, potentially as the earliest branching member within nudiform loricates, which would be informative for investigating the evolution of the nudiform lifestyle. Our findings are inconsistent with a strict classification based on currently defined aspects of lorica morphology and support the usage of genetic data as primary criterion for genus-level taxonomic assignment.

## INTRODUCTION

Choanoflagellates are a clade of heterotrophic, bacterivorous flagellates characterized by a distinct body plan consisting of a cell body, a collar of microvilli, and a single flagellum. They use these structures to capture and ingest prey: the flagellum beats to generate a water current that draws bacteria towards the collar, where the bacteria become trapped on the microvilli and are then transported to the cell body for phagocytosis (Leadbeater, 2015; Richter & Nitsche, 2016) .

Choanoflagellates share a clear morphological similarity with some animal cell types, such as sponge choanocytes (Brunet & King, 2017), which together with their key phylogenetic position as the sister clade to Metazoa, makes them a clear target to study the emergence of Metazoa, animal cell biology and the evolution of animal multicellularity and temporal and spatial cell differentiation (Ros-Rocher et al., 2021). Choanoflagellates are globally present in all aquatic environments, deeply influencing microbial food webs and the ecology of our planet (Leadbeater, 2015) .

Choanoflagellate phylogeny is divided mainly in two orders: Craspedida and Acanthoecida (Nitsche et al., 2011); both orders contain marine and freshwater species. The craspedids have representatives with three different types of extracellular structures, the glycocalyx, a fine and flexible mucilaginous cover; the theca: a more rigid organic structure, also subdivided by shape; and choanoflagellates without extracellular structure, referred to as naked choanoflagellates. The acanthoecids are a monophyletic group which possess another extracellular structure: the lorica, which is a silica-based basket-like structure composed of costal strips. Within Acanthoecida, loricates are divided in two types depending on their division mode: nudiforms, which undergo a brief “naked” motile stage after cell division before constructing their own lorica, and tectiforms, in which the mother cell assembles the daughter cell’s lorica prior to cytokinesis and a “naked” motile dispersal stage is not present in typical culture conditions. The tectiform choanoflagellates are assigned to the Stephanoecidae family, while nudiforms are assigned to the family Acanthoecidae (Nitsche et al., 2011), although nudiforms have recently been found to branch within tectiforms (Carr & Leadbeater, 2022) .

In this study, we characterize cell cultures of two choanoflagellate strains, allowing us to explore the relationship between morphological and genetic variation within loricate (Thomsen et al., 1997; Thomsen & Larsen, 1992; Thomsen & Østergaard, 2017)choanoflagellates. Loricate choanoflagellates have traditionally been classified based on the morphology of their defining characteristic, the lorica, but our study provides evidence that morphology alone may be insufficient for phylogenetic assignment. Discrepancies when comparing the relationships among species based on either genetic or morphological data give rise to questions such as: Can a single loricate species exhibit multiple morphologies? Conversely, can two different species share the same lorica morphology? Or could two different species share the same 18S rRNA? Our findings of unexpected features when comparing both strains to their nearest genetic relatives highlights the need for further revision of the standard procedures applied to interpreting morphological evolution within loricate choanoflagellates.

## RESULTS

### Phylogenetic position of two loricate choanoflagellates

We isolated and brought into eukaryotic monoculture two strains that we identified as loricate choanoflagellates, BEAP0094 and BEAP0360 (the latter also known under the strain names SIOpierAcanth1 and 10tr). To place these two cultures within choanoflagellate diversity and to help interpret the evolutionary patterns of their morphology and behavior, we selected 23 publicly available choanoflagellate proteomes (8 from loricates, including BEAP0360, which was sequenced as part of the Moore Microbial Eukaryote Transcriptome Sequencing Project) (Keeling et al., 2014; Richter et al., 2022). We sequenced and assembled the transcriptome of BEAP0094 and produced a predicted proteome. Next, we created a phylogenomic dataset with PhyloFisher (Tice et al., 2021) followed by tree reconstruction with IQ-TREE (Nguyen et al., 2015) (Figure 1). The resulting tree topology is consistent with previous studies based on six to nine genes typically used for phylogenetic reconstruction in choanoflagellates (Carr et al., 2017; Carr & Leadbeater, 2022; Ginés-Rivas & Carr, 2025; Hake et al., 2024), with the exception of the branching order among the three nudiforms *Acanthoeca spectabilis*, *Savillea parva*, and *Helgoeca nana*, which differs from Ginés-Rivas & Carr. This discrepancy could be due to the short branch lengths separating these three taxa, as the nodes separating them receive less than full support in Ginés-Rivas & Carr and in our 9-gene tree (see below). In our phylogenomic reconstruction, all nodes within loricate choanoflagellates (Acanthoecida) show full bootstrap support. BEAP0094 branches inside the tectiforms (Stephanoecidae), sister to *Diaphanoeca grandis*. BEAP0360 branches as the sister to the clade composed of the 3 nudiforms (Acanthoecidae) present in the tree.

**Figure 1.**
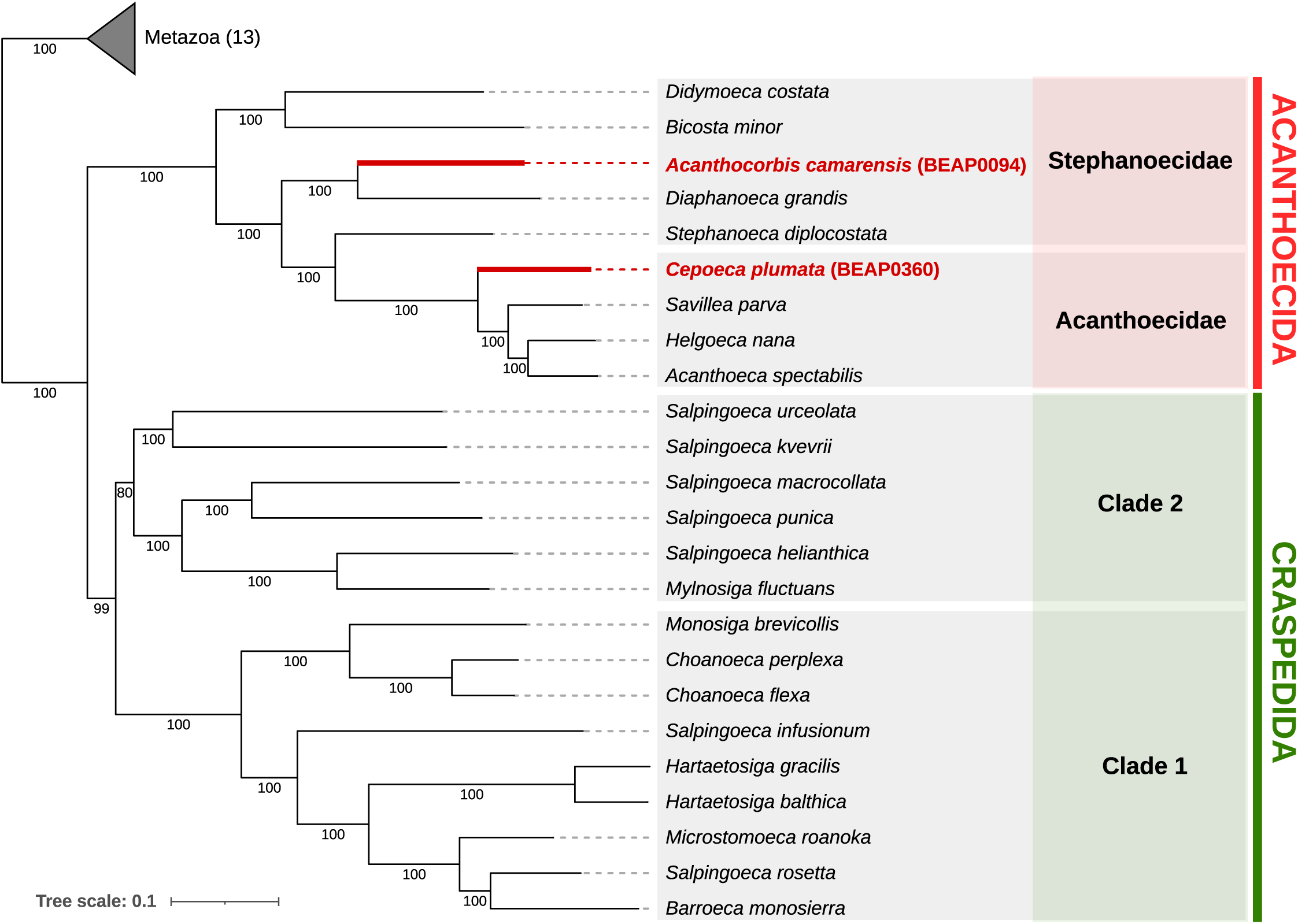
BEAP0094 and BEAP0360 phylogenetic position within Choanoflagellatea. Phylogenetic tree of Choanoflagellatea diversity based on 230 reconstructed with Phylofisher and IQ-TREE using the LG+F +Rl0 model of nee evolution. Node support reflects 1000 bootstraps. BEAP0094 branches with pport within Stephanoecidae, as sister to *Diaphanoeca grandis,* while BEAP0360 hes with full support as sister to currently sequenced Acanthoecidae, and is lered part of this family due to its nudiform mode of reproduction. Number of species included in Metazoa outgroup is indicated in parenthesis. The tree was ally rooted between Choanoflagellatea and Metazoa.

In order to corroborate the phylogenetic placement of these two strains, we decided to build a tree with a larger sampling of choanoflagellate species, focusing on loricates. Unfortunately, for many species only limited sets of genes were publicly available; therefore, we built a 9-gene phylogenetic tree, combining nucleotide (18S and 28S) and amino acid (actin, alpha tubulin, beta tubulin, EFL, EF1A, HSP70 and HSP90) genes, following the approach established in (Carr & Leadbeater, 2022), with available loricate choanoflagellates, providing a higher resolution inside Acanthoecida (Figure 2). All branches within Acanthoecida in the resulting 9-gene phylogeny are consistent with the topology of the phylogenomic tree. The increased number of loricate species in the 9-gene tree allowed us to interpret the phylogenetic position of our two strains at increased resolution. BEAP0094 branches inside the tectiform family Stephanoecidae, sister to *Pseudostephanoeca paucicostata*, whereas BEAP0360 branches sister to all currently sequenced nudiforms. All loricate genera are monophyletic, with the exception of *Stephanoeca*, corroborating previous studies that identified it as a genus in need of revision (Nitsche et al., 2011) .

**Figure 2.**
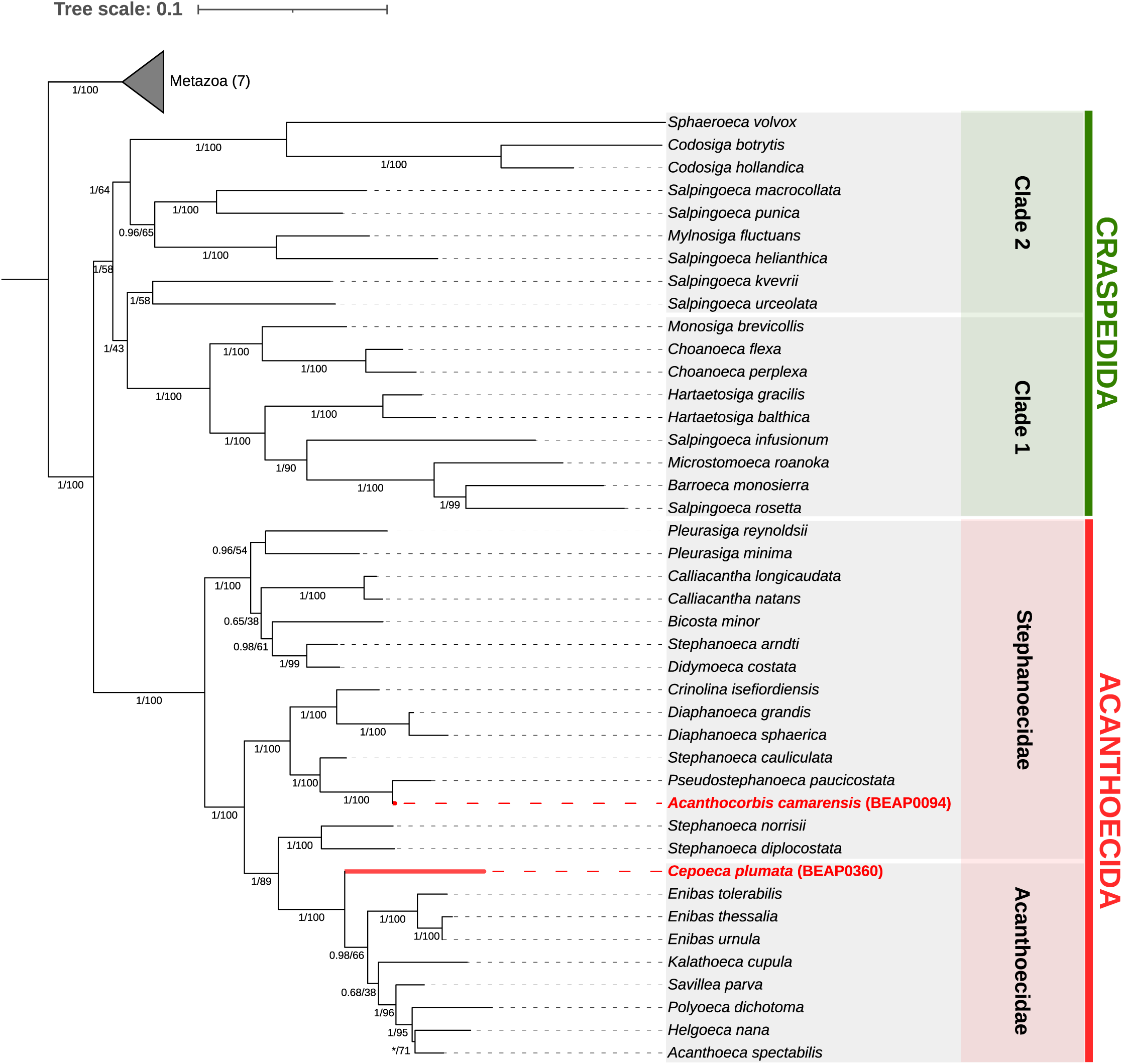
Phylogeny of Choanoflagellatea based on nucleotide (18S and 28S and amino acid (actin, β-tubulin, EFlA, HSP70, α-tubulin, EFL, HSP90, and 1ene sequences. The topology was reconstructed using RAxML, employing the bstitution model for nucleotide genes and the WAG model for amino acid genes, with 1000 ultrafast bootstrap replicates. Bayesian inference was conducted using MrBayes to confirm the maximum likelihood topology. Amino acid partitions were analyzed under a amino acid substitution model (aamodelpr=mixed) with gamma-distributed rate variation, while nucleotide partitions used a GTR+l+Γ model (nst=6, rates=invgamma) ffiable substitution rates across partitions. Two independent MCMC analyses of 1,000,000 generations were run, each with four chains and sampling every 100 generations; convergence diagnostics were checked every 10,000 generations, and the first 25% of samples were discarded as burn-in. Node support values are shown as Jr probabilities (left, MrBayes) and bootstrap percentages (right, RAxML). An c (*) indicates node not found in MrBayes consensus tree, where *Helogoeca nana* is inferred as sister to the *Polyoeca dichotoma* + *Acanthoeca spectabilis* clade with a posterior probability of 0.56.

In the 9-gene tree, the BEAP0094 18S rRNA gene sequence is very closely related to that of *Pseudostephanoeca paucicostata* (Schiwitza et al., 2022) (Figure 2), a species that is morphologically distinct, as our isolate does not possess a *Stephanoeca*-like anterior ring (discussed below). This observation implies that either a single loricate species may exhibit multiple morphologies, or that two different morphological species share a very similar 18S rRNA gene sequence.

### Morphological characterization of two strains of loricate choanoflagellates

BEAP0094 is a solitary loricate choanoflagellate. Its cell body is ∼5.2 μm long (n = 3 for all measurements), ∼3.6 μm wide, and, as is typical for loricate choanoflagellates, is located inside the posterior chamber. The microvillar collar is ∼4.4 μm long, while the flagellum is ∼7.4 μm long. The obpyriform-shaped lorica is composed of 4-6 transverse costae and 14-16 anterior spines, without the presence of an anterior ring. The posterior transverse costae form a flange that separates the anterior (width ∼10.6 μm) and posterior (width ∼5.4 μm) chamber of the lorica. The lorica is ∼12.5 μm long (including spines) and the costal strips are ∼5.6 μm long (Figure 3). The division mode is tectiform, as expected by its phylogenetic position (Supplementary Figure 1, Supplementary video 1). This morphology is consistent with *Acanthocorbis camarensis* (Hara et al., 1996); a more in-depth justification is provided below in the Discussion.

**Figure 3.**
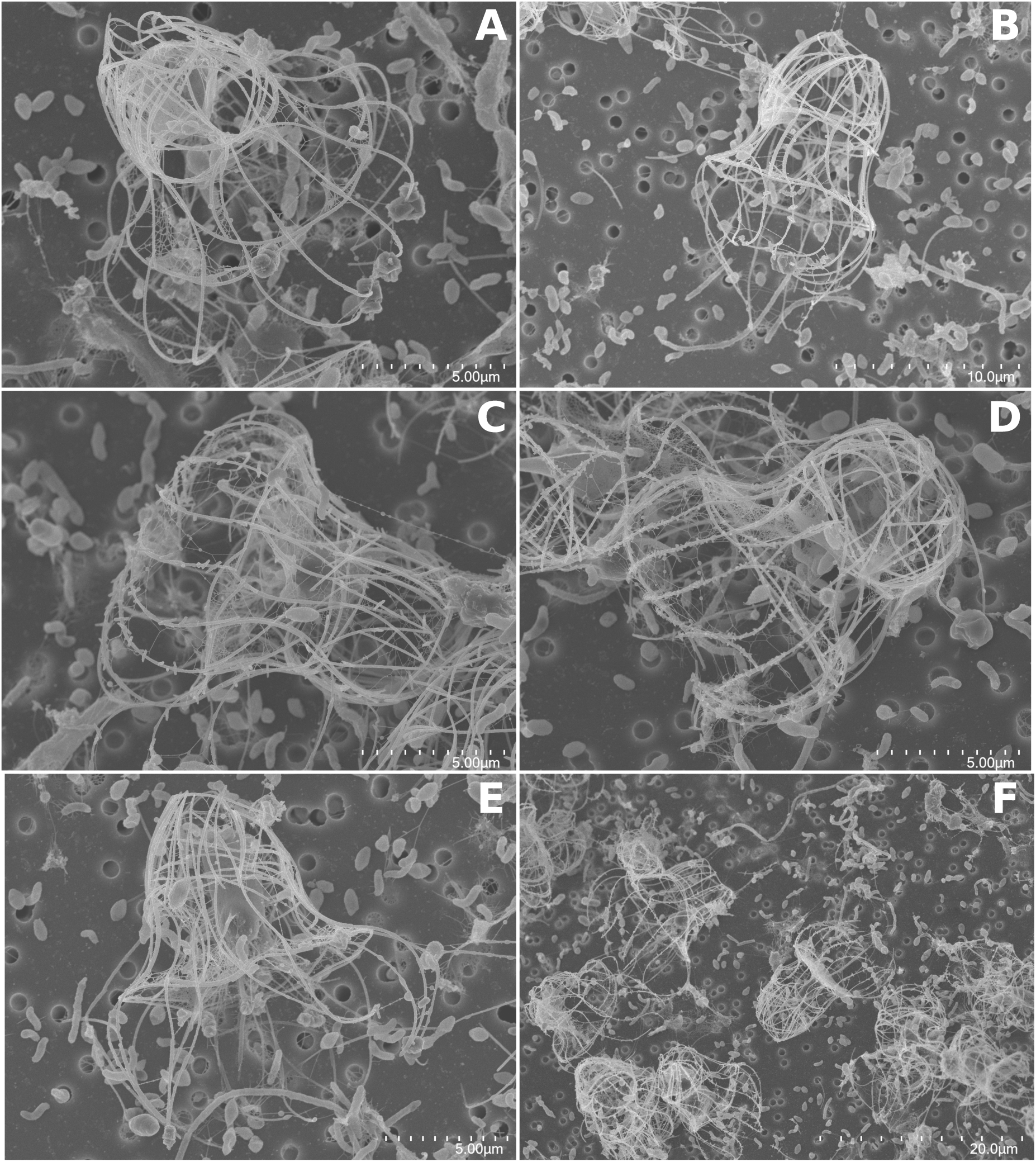
Scanning electron microscopy of BEAP0094. **A to E**. Single cells with multiple morphologically diverse prey bacteria. F. Multiple cells with multiple morphologgically diverse prey bacteria. Scale bar sizes indicated in each panel.

BEAP0360 is a solitary loricate choanoflagellate with a stalk. Its cell body is ∼4.6 μm long (n = 3 for all measurements), ∼3.2 μm wide, and located inside the posterior chamber. The microvilli collar is ∼4.3 μm long, while the flagellum is ∼7.9 μm long. The champagne glass-shaped lorica is composed of 2-3 transverse costae and 8-10 anterior spines, with the presence of an anterior ring. The posterior costae twist in a way that generates a long and posterior protuberance and separate the anterior (width ∼7 μm) and posterior (width ∼4.6 μm) chambers of the lorica. The lorica is ∼16.8 μm long (including spines and anterior structures) and the coastal strips are ∼4.7 μm long (Figure 4). This lorica morphology is consistent with that of *Stephanoeca cauliculata* (Leadbeater, 1980).

**Figure 4.**
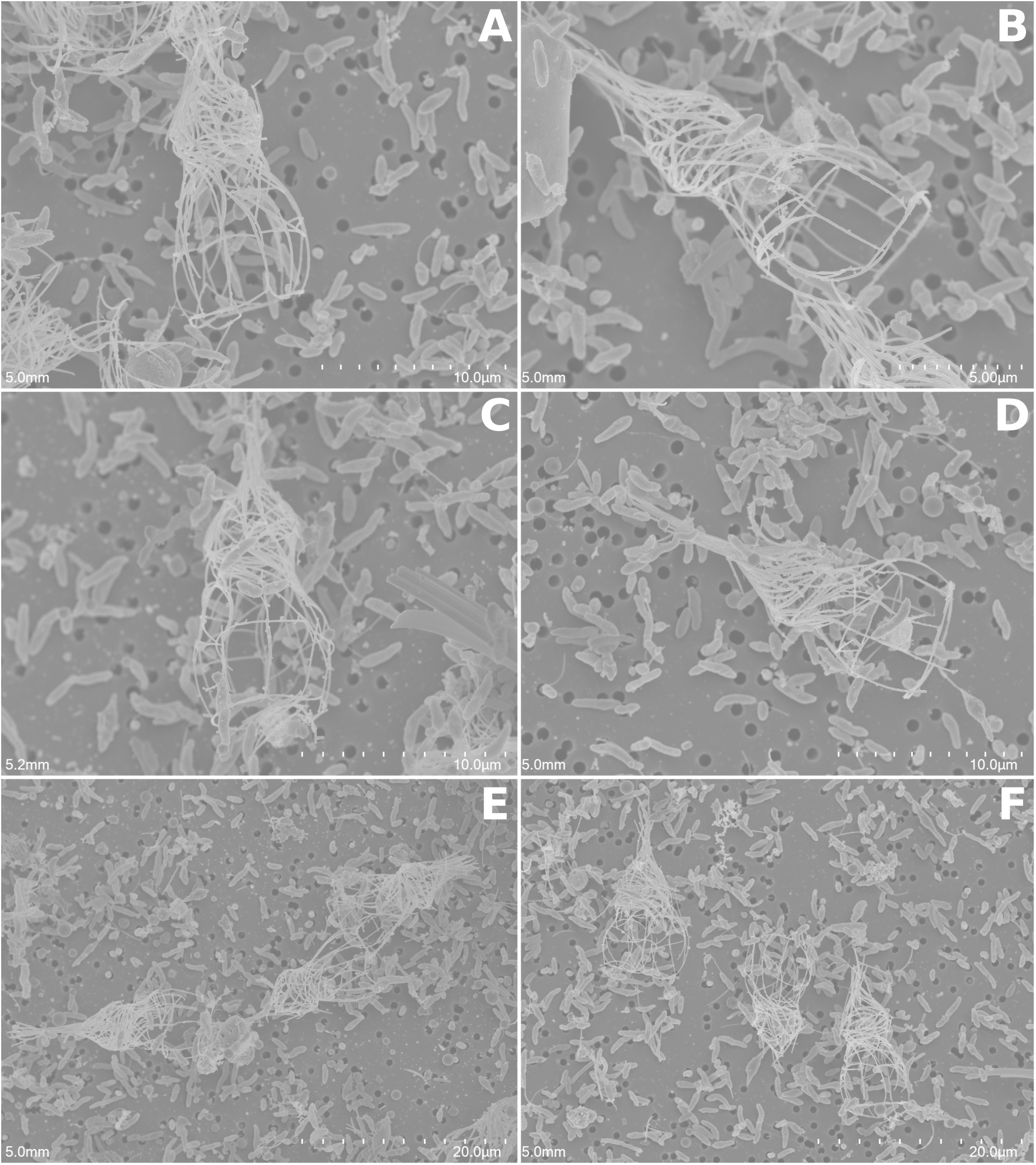
Scanning electron microscopy of BEAP0360. **A to D**. Single cells with multiple morphologically diverse prey bacteria. E, F. Multiple cells with multiple morphologgically diverse prey bacteria. Scale bar sizes indicated in each panel.

Given the phylogenetic position of BEAP0360 as sister to all currently sequenced loricate species described as nudiforms, we decided to explore its division mode by time-lapse microscopy. We found that its division can be nudiform-like (Supplementary Figure 2, Supplementary video 2), and through extensive time-lapse microscopy, we never observed tectiform-like division (∼25 8h time lapses). Our observations are therefore consistent with a nudiform division mode.

BEAP0360 shows morphological features consistent with *Stephanoeca cauliculata*, yet its 18S rRNA sequence diverges from the only reported sequence for that species (Nitsche et al., 2011). We also present observations of its nudiform mode of division, which would be inconsistent with its assignment to *Stephanoeca*, a tectiform genus. This raises the possibility that two different species may share convergently evolved and similar lorica morphologies.

## DISCUSSION

In this study, we analyzed cultures of two loricate choanoflagellates, BEAP0094 and BEAP0360, to characterize their morphology and phylogenetic placement. In the course of our analyses, we encountered two disagreements between morphological and phylogenetic relationships.

First, our isolate BEAP0094 very closely matched (1.46 substitutions per 100 sites in the alignment used to generate Figure 2) the 18S ribosomal gene of *Pseudostephanoeca paucicostata*, but its morphology clearly differed, mainly due to the absence of the anterior ring that is characteristic of all *Stephanoeca* species (Ellis, 1930; Thomsen & Østergaard, 2019) Instead, BEAP0094 more closely matched the morphology *Acanthocorbis* (Hara & Takahashi, 1984), a genus described before sequencing the 18S ribosomal gene was a standard practice.

We observed that for BEAP0094, nearly all the 18S differences that separate it from *P. paucicostata* are in the *P. paucicostata* sequence (Figure 2), suggesting the possibility that *P. paucicostata* experienced rapidly accelerated evolution following its divergence with BEAP0094 (it is also formally possible that errors may have been introduced during its sequencing process). Their close relationship also raised the possibility of the existence of multiple lorica morphologies within the same (or very closely related) species or to the existence of multiple morphological species sharing a nearly identical 18S ribosomal gene.

We were never able to observe, over 3 years of culturing, any loricae of BEAP0094 featuring an anterior ring. Therefore, either the various culture conditions that we used favor the construction of only a single lorica type without an anterior ring, and the species would be capable of producing a *Stephanoeca*-type lorica in appropriate conditions, or instead, we have identified a case in which two closely genetically related species produce different lorica types, one *Acanthocorbis*-like (BEAP0094) and the other *Stephanoeca*-like (*Pseudostephanoeca paucicostata*). The former would be consistent with previous observations in fixed environmental samples, in which *Acanthocorbis*-like and *Stephanoeca*-like were found in close physical contact, in a manner suggestive of a mother/daughter lorica relationship (Thomsen & Østergaard, 2017).

Regarding the identity of BEAP0094 within the genus *Acanthocorbis*, based on the descriptions of *A. apoda* (Leadbeater, 1972) and *A. camarensis* (Hara et al., 1996), which are differentiated by the number of transverse costae in the posterior chamber and in the spacing of the first and second transverse costae, we identified BEAP0094 as *A. camarensis*. Specifically, *A. camarensis* is characterized by having more than three transverse costae in the posterior chamber, whereas *A. apoda* has only one to three. Additionally, in *A. camarensis*, the first and second transverse costae are separated by less than one longitudinal coastal strip, while in *A. apoda* the separation is approximately one and a half longitudinal coastal strips. Therefore, BEAP0094 matches the *A. camarensis* description, with more than three transverse costae and a separation of less than one longitudinal coastal strip between the first and second transverse costae. Nonetheless, the presence of relatively small variations in the number of anterior spines and transverse costae, lead us to consider the possibility that *A. camarensis* and *A. apoda* could be ecotypes of the same species, as their differentiating traits might be explained by environmental effects. If this were the case, the identification of BEAP0094 would need to be changed to *A. apoda*, as its earlier description would take precedence.

The second discrepancy that we encountered was that our isolate BEAP0360 presented a morphological match to *Stephanoeca cauliculata*, but its 18S ribosomal sequence is clearly different (Figure 2). This raises the possibility of multiple genetic species sharing the same lorica morphology, although it is also formally possible that the conflict was caused by a misidentification during the sequencing of the *Stephanoeca cauliculata* 18S sequence, as this species could not be successfully cultured and instead its sequence was produced via single-cell PCR.

The strain BEAP0360 possesses a key phylogenetic placement as, potentially, the earliest branching nudiform choanoflagellate (Acanthoecidae; Figure 1, Figure 2). Our observations indicated that its division mode is nudiform-like (Supplementary video 2), meaning that it should be designated as the earliest-branching choanoflagellate inside Acanthoecidae. Therefore, although this species morphologically matches the *Stephanoeca* genus, it cannot be considered to be a *Stephanoeca* species, as this genus, and all other members of Stephanoecidae reproduce by tectiform division (Nitsche et al., 2011). Supported by its phylogenetic placement as sister to sequenced nudiforms (Acanthoecidae) species, we propose a new nudiform genus, *Cepoeca* within the family Acanthoecidae, and a new species, *Cepoeca plumata*.

The identification of *Cepoeca* has implications for the evolution of the nudiform condition. As a sister to described and sequenced nudiform genera, it provides a comparison point with the rest of Acanthoecidae. In addition, its morphology and behavior can be interpreted to understand the evolution of nudiforms from what was likely a tectiform common ancestor. Its lorica morphology displays the signature anterior ring present in the tectiform genus *Stephanoeca*, implying that this feature can be present within both nudiforms and tectiforms, and that its presence does not necessarily identify a species as tectiform. Indeed, *Enibas*, another recently described nudiform genus, is also characterized by a lorica morphology with *Stephanoeca*-like features (Leadbeater & Carr, 2022; Schiwitza et al., 2019; Schiwitza & Nitsche, 2021; Schiwitza & Thomsen, 2022). In our phylogenetic trees, (Figure 1, Figure 2), two members of the genus *Stephanoeca* are the most closely related to the clade containing *Cepoeca* and previously described nudiforms: *S. diplocostata* and *S. norrisii*. Of these two species, only *S. diplocostata* has been investigated in a laboratory setting, where it was found that, although the cells typically reproduce by tectiform division, when completely deprived of silicon in their growth medium, they are capable of nudiform division and accumulation of costal strips in a nudiform-like manner when silicon is restored to the medium (Leadbeater, 1989; Leadbeater & Carr, 2022). Similar observations in the distantly-related loricate species *Diaphanoeca grandis*, in which a subset of cells were capable of nudiform-like costal strip accumulation after silicon was restored to a silicon-free medium (Leadbeater, 2015), combined with the emerging consensus that nudiform species emerged from within tectiforms (Figure 1, Figure 2; Carr & Leadbeater, 2022) support the hypothesis that tectiform division was ancestral in loricate choanoflagellates and subsequently lost in the nudiform clade. Nonetheless, it remains to be investigated which aspects of nudiform division and lorica construction are shared by all loricate species, shared by *S. diplocostata* and the nudiforms, or nudiform-specific. Future comparisons of the gene content of *C. plumata* with tectiform and nudiform species, coupled with the development of gene manipulation techniques in loricates, could greatly contribute to understanding the genetic basis of the nudiform lifestyle.

Our findings related to our two isolates are inconsistent with the assignment of species to genera based on morphological traits alone. *Acanthocorbis camarensis* BEAP0094 does not morphologically resemble its closest sequenced tectiform relatives, as it does not possess the *Stephanoeca*-like anterior ring. Conversely, *Cepoeca plumata* BEAP0360 is a nudiform whose anterior ring morphologically matches the lorica characteristics of the tectiform genus *Stephanoeca*, but it is instead more closely related to nudiform loricates. Nevertheless, the genera *Calliacantha, Pleurasiga* and *Diaphanoeca*, which were described prior to the adoption of DNA sequencing, together with the more recently described genus *Enibas* are all monophyletic in both our phylogenomic and 9-gene phylogenies, hinting towards some clear aspects of morphological conservation. In fact, it may be that the presence of an anterior ring will need to be set aside as an identifying characteristic of loricate genera, based on our findings and on the presence of species with *Stephanoeca*-like morphologies throughout the loricate choanoflagellate tree. Nonetheless, we can be confident that not all acanthoecid genera have morphologically conserved loricae, suggesting that future genus-level taxonomic assignment should rely primarily on genetic data, as morphology-based classification can be misleading.

## TAXONOMIC SUMMARY

### Eukaryota, Amorphea, Obazoa, Opisthokonta, Holozoa, Choanozoa, Choanoflagellatea, Acanthoecida, Acanthoecidae

*Cepoeca* Gàlvez-Morante and Richter, n. gen.

Description: Solitary nudiform choanoflagellate. Cell body located inside the posterior chamber. Microvillar collar surrounding a single flagellum. Champagne glass-shaped lorica composed of multiple transverse costae and anterior spines, with an anterior ring.

Etymology: Named for the Latin word “cepa”, meaning “onion”, in reference to the onion dome architectural element found in buildings with towers throughout Central and South Asia, the Middle East and Eastern and Western Europe. The stalk of the lorica resembles the spire of an onion dome, the posterior chamber resembles the onion dome itself, and the anterior chamber resembles its lantern. *Cepoeca* is feminine.

Type species: *Cepoeca plumata* (see below).

Zoobank registration: Described under the Zoological Code; Zoobank registration will be performed following peer review.

*Cepoeca plumata* Gàlvez-Morante and Richter, n. sp.

Description: *Cepoeca plumata* is a solitary nudiform choanoflagellate with stalk. Its cell body is ∼4.58 μm long (n = 3 for all measurements), ∼3.22 μm wide, and located inside the posterior chamber. The microvilli collar is ∼4.3 μm long, while the flagellum is ∼7.85 μm long. The champagne glass-shaped lorica is composed by 2-3 transverse costae and 8-10 anterior spines, with the presence of an anterior ring. The posterior costae twist in a way that generate a long and posterior protuberance and separate the anterior (width ∼7 μm) and posterior (width ∼4.6 μm) chambers of the lorica. The lorica is ∼16.78 μm long (including spines and anterior structures) and the coastal strips are ∼4.7 μm long.

Etymology: The Latin term “plumata” refers to its appearance, as its posterior chamber looks like a feathered badminton shuttlecock.

Type locality: Coastal water in the United States, in Scripps Coastal Reserve (32° 52’ 1.2” N / 117° 15’ 18.0” W).

Type material: The name-bearing type (an hapantotype) is an SEM stub deposited in the Marine Biological Reference Collections (CBMR) at the Institut de Ciències del Mar (ICM-CSIC, Barcelona, Spain) under the catalog/accession number ICMCBMR000697 (Guerrero et al., 2023). This material also contains uncharacterized prokaryote species, which do not form part of the hapantotype.

Gene sequence: The 18S ribosomal gene sequence of strain BEAP0360 (also known by the strain names SIOpierAcanth1 and 10tr) is available in Genbank with the identifier KM516200.

Cell culture: A culture containing BEAP0360 and mixed bacterial species is publicly available and has been deposited in the Culture Collection of Algae and Protozoa as CCAP 2954/1.

Zoobank registration: Described under the Zoological Code; Zoobank registration will be performed following peer review.

This publication (work) will be registered with Zoobank following peer review.

## METHODS

### Isolation and culture

BEAP0094 was isolated from the coast of Spain, in Blanes Bay (41°40’ N / 2°48’ E) in May, 2022 through the dilution of surface marine water in RS medium (1 NM : 50 NNM concentration). RS medium is a fully defined medium designed to grow a wide range of marine protists. It is composed of two components: the nutrient medium (NM) and the non-nutrient medium (NNM) (https://mediadive.dsmz.de/medium/P4, https://mediadive.dsmz.de/medium/P5) (Sigona *et al*., in preparation).

BEAP0094 was maintained in 25 cm^2^ flasks with a total volume of 10 mL, with a split frequency of 1 split per 1.5 weeks at 18 °C, in the presence of light. The splits were performed by adding 9 mL of R/S medium (1 NM : 50 NNM concentration) and 1 mL of the previously growing culture, without scraping.

BEAP0360 was isolated from the coast of the United States, in Scripps Coastal Reserve (32° 52’ 1.2” N / 117° 15’ 18.0” W) during enrichments for grazers of the cyanobacterium *Synechococcus*.

BEAP0360 was maintained in 25 cm^2^ flasks with a total volume of 10 mL, with a split frequency of 1 split per 1.5 weeks at 18 °C, in the presence of light. The splits were performed by adding 9 mL of F/4 medium (1:2 dilution of F/2 medium; Guillard, 1975) and 1 mL of the previously growing culture, without scraping. The cultures include *Synechococcus*.

### BEAP0094: 18S rRNA gene sequencing

For BEAP0094, a sample of 50 mL (two 75 cm^2^ flasks with a total volume of 25 mL) was used for 18S cloning. The cells were centrifuged for 20 minutes at 13000 x g and 4 °C. DNA was extracted using the DNeasy PowerSoil Pro kit (QIAGEN) and the 18S rRNA gene was amplified by PCR with universal eukaryotic primers (CTCAARGAYTAAGCCATGCA - CCTTCYGCAGGTTCACCTAC) (López-García et al., 2003; Vaulot et al., 2022). Amplification products were purified using NZYGelpure kit (NZYTech), cloned using the TOPO-TA Cloning kit (Invitrogen) and transformed into *E. coli* cells following LacZα-complementation. Positive clones were selected and amplified by PCR with vector-specific primers M13F-M13R (GTAAAACGACGGCCAGT - CAGGAAACAGCTATGAC). Sequencing was performed by Eurofins genomics using Sanger sequencing. The resulting sequences were base called using phred (Ewing et al., 1998) with the parameters ‘-trim_alt “” -trim_cutoff 0.01’ and assembled with phrap (de la Bastide & McCombie, 2007) with the parameter ‘-repeat_stringency 0.4’ and the consensus sequence was exported with consed (Gordon & Green, 2013). Assembled sequences were identified through BLAST versus EukRibo (Berney et al., 2022). This sequence was used in phylogenetic trees and is identical to the 18S sequence contained in the ribosomal operon deposited in GenBank with the identifier PV791352.

For BEAP0360, the sequence used in phylogenetic trees was obtained from the publicly available metadata associated with dataset EP00036 distributed as part of the EukProt v3 database (Richter et al., 2022). As described in the EukProt v3 metadata, the complete 18S sequence was extracted from contig CAMNT_0025056341 from the MMETSP0105 transcriptome assembly of the Marine Microbial Eukaryote Transcriptome Sequencing Project (Keeling et al., 2014) .

### BEAP0094: RNA extraction

In order to extract and sequence total RNA we used a similar experimental approach as the one described by Richter and colleagues (Richter et al., 2018). Prior to RNA isolation, both strains were grown in large batches using two 75 cm^2^ Rectangular Canted Neck Cell Culture Flasks with Vented Caps (353136, Corning Life Sciences), each containing 50 ml of medium.

We isolated total RNA from both strains by using the RNAqueous kit (AM1912, ThermoFisher Scientific). RNA concentration was measured using a NanoDrop One/One C UV-Vis Microvolume Spectrophotometer (ThermoFisher Scientific). We digested genomic DNA using the TURBO DNA-free kit (AM1907, ThermoFisher Scientific) according to the manufacturer’s instructions and removed DNase using DNase Inactivation Reagent. The quality evaluation of extracted total RNA, the library preparation, and the sequencing were carried out at the CRG Genomics Core Facility in Barcelona. The quality of extracted total RNA was assessed by Bioanalyzer 2100 RNA Pico chips (Agilent Technologies). To prepare stranded libraries, the NEBNext Ultra II Directional RNA Library Prep Kit Total RNA was used, applying double the normal rounds of PolyA selection prior to cDNA synthesis to reduce the excess of bacterial RNA. Finally, the RNA was sequenced by using Illumina NextSeq 2000 (2x150 bp).

### BEAP0094: *De novo* transcriptome assembly and proteome generation

The quality assessment of the sequenced RNA was performed using FastQC v0.11.9 (Andrews S., 2010), while evaluating the purity of the sequences using phyloFlash v3.4 (Gruber-Vodicka et al., 2020), a tool designed to assess the 16S/18S rRNA gene taxonomic composition of metagenomic datasets. The elimination of artifacts and adapter sequences was achieved via fastp v0.23.2 (Chen et al., 2018) with the following parameters: --low_complexity_filter, --cut_front, --cut_tail, --cut_right, --cut_front_window_size 1 --cut_tail_window_size 1, --cut_right_window_size 4, --cut_mean_quality 5, --trim_front1 12 --trim_front2 12, --trim_poly_g, --trim_poly_x --adapter_sequence=AGATCGGAAGAGCACACGTCTGAACTCCAGTCA --adapter_sequence_r2=AGATCGGAAGAGCGTCGTGTAGGGAAAGAGTGT. *De novo* transcriptome assembly was performed using Trinity v2.14.0 (Grabherr et al., 2011) with parameter --SS_lib_type RF.

We decontaminated the newly assembled transcriptome by blastn against a set of potential contaminant genomes (species with a 16S/18S detected by phyloFlash or Trinity; Supplementary file 1) and removing matching contigs with a percentage identity of ≥ 98% and match length > 100. WinstonCleaner with default parameters was also used to remove cross-contamination between samples sequenced in the same run (https://github.com/kolecko007/WinstonCleaner). To examine the completeness of our transcriptomes, we searched for the presence of a set of conserved, single-copy orthologous genes by employing BUSCO v5.3.2 with the Eukaryota odb10 dataset (Simão et al., 2015).

### Phylogenetic tree inference

The transcriptomes and predicted proteomes of BEAP0094 and BEAP0360 underwent PhyloFisher’s protocol for phylogenomic assembly (Tice et al., 2021), together with 22 added choanoflagellate species from the EukProt v3 database (Richter et al., 2022), following the standard PhyloFisher instructions (https://thebrownlab.github.io/phylofisher-pages/). The PhyloFisher protocol includes the collection of homologs for the input species with HMMER (Mistry et al., 2013); construction of gene trees: aligned using MAFFT (Katoh & Standley, 2013), filtered with PREQUAL (Whelan et al., 2018) + DIVVIER (Ali et al., 2019), trimmed with BMGE (Criscuolo & Gribaldo, 2010) and trimAl (Capella-Gutiérrez et al., 2009) and built with RAxML (Stamatakis, 2014); manual inspection of gene trees to identify and remove paralogs and contaminants using ParaSorter (included with PhyloFisher); followed by re-alignment; trimming using the same procedure; and concatenation of the selected orthologs into a supermatrix. The final maximum likelihood tree was generated with IQ-TREE, which with ModelFinder selected the LG+F+R10 model, with 1000 ultrafast bootstraps (Nguyen et al., 2015).

We also performed phylogenetic tree inference using the 18S, 28S, actin, alpha tubulin, beta tubulin, EF1A, EFL, HSP70 and HSP90 genes, and conducted a deeper sampling of Choanoflagellatea to validate the PhyloFisher results. To enlarge our taxon sampling, we included sequences from previous works (Carr & Leadbeater, 2022; Frank et al., 2017; Hake et al., 2024; Leadbeater & Carr, 2022; Richter et al., 2018; Schiwitza et al., 2019, 2022, 2022; Schiwitza & Nitsche, 2021; Schiwitza & Thomsen, 2022), either from the data included as part of the publications or from GenBank (*Barroeca monosierra* 18S MW838180.1, EFL MW979373.1 and Hsp90 MW979374.1; *Kalathoeca cupula* 18S OP764483.1 and 28S OP771620.1). We preferentially selected *Salpingoeca rosetta* and *Monosiga brevicollis* sequences to blastn against our two input transcriptomes to recover these 9 genes, keeping the sequences that were the best hit in both cases. Due to the lack of sequences from these species for the EF1A gene, we used as query the sequences of *Salpingoeca helianthica* and *Mylnosiga fluctuans* .

Ribosomal genes were aligned with fsa and trimmed with trimal (-gt 0.5), while non-ribosomal genes were translated to protein sequences. To identify the correct reading frames for protein sequences, we translated our transcriptomes across all 3 open reading frames (ORFs) (1_find_rosetta_genes.sh). We then manually blasted *S. rosetta* results against the NCBI database to determine the correct amino acid sequence for each gene (except for EF1A, for which we used the *M. musculus* ortholog, as no choanoflagellate proteins derived from genomic sequences are currently available). Using this information, we blasted the *S. rosetta* sequences against all translated proteins to confirm and retain only the best hit per species, which we aligned with fsa (2_select_correct_orfs.sh). The resulting alignments were manually trimmed using the UGENE graphical interface (Okonechnikov et al., 2012) .

Gene trees were inferred with RAxML v.8.2.12 (Stamatakis, 2014) for both ribosomal and non-ribosomal genes, using the GTRCAT model for nucleotide data and the PROTCATDAYHOFF model for amino acid data. We then used PhyKIT (Steenwyk et al., 2024) to concatenate alignments across all genes, generating a combined matrix.

The concatenated matrix was employed to reconstruct a maximum-likelihood tree in RAxML using both nucleotide and amino acid data. The GTR substitution matrix was applied to nucleotide genes and the WAG substitution matrix was applied to amino acid genes, with 1000 ultrafast bootstraps. We also used MrBayes v3.2.6 (Ronquist et al., 2012) to reconstruct a Bayesian inference phylogeny to compare with the maximum likelihood results. For protein sequences, we used a mixed amino acid substitution model (aamodelpr=mixed) with gamma-distributed rate variation across sites. The 18S and 28S genes were analyzed under the GTR+I+Γ model (nst=6, rates=invgamma) with variable substitution rates across partitions, following Carr et al., 2022. Two independent Markov Chain Monte Carlo (MCMC) analyses were run in parallel for 1,000,000 generations, each using four chains and sampling every 100 generations. Convergence diagnostics were assessed every 10,000 generations. The first 25% of samples were discarded as burn-in.

### Scanning Electron Microscopy

Cells were fixed with 2.5% glutaraldehyde for 3 hours at room temperature and then seeded on membranes with 0.8 μm pores (WHA10417301, MERCK Chemicals and Life Science) via filtering. The samples were dehydrated through a graded ethanol series and dried by critical point with liquid carbon dioxide in a Leica EM CPD300 unit (Leica Microsystems, Austria). The dried filters were mounted on stubs with colloidal silver and then were sputter-coated with gold in a Q150R S (Quorum Technologies, Ltd.) and observed with a Hitachi SU8600 field emission scanning electron microscope (Hitachi High Technologies Co., Ltd., Japan) in the Electron Microscopy Service of the Institute of Marine Science (ICM-CSIC), Barcelona.

## DATA, CODE AND MATERIALS AVAILABILITY

Data: BEAP0094 transcriptome reads and assembled contigs are available in GenBank under BioProject accession number PRJNA1276494. The predicted proteins (as well as the contigs from the assembly) are available on FigShare at DOI 10.6084/m9.figshare.29503316. The transcriptome assembly available on GenBank differs from the one on FigShare due to the automatic decontamination tools used by GenBank, which we believe have erroneously removed a number of valid contigs. As a result, we recommend using our FigShare assembly, which retains these sequences, over the GenBank versions. The ribosomal operon sequence (18S, ITS1, 5.8S, ITS2, 28S) sequence of BEAP0094 is available with GenBank accession number PV791352. BEAP0360 transcriptome contigs and predicted proteins were produced as part of the Moore Microbial Eukaryote Transcriptome Sequencing Project and are available on FigShare with the DOI 10.6084/m9.figshare.12410606. The same set of predicted proteins is also available via EukProt v3 with the identifier EP00036. The 18S ribosomal sequence is available with GenBank identifier KM516200. Phylogenetic trees, input sequences and alignments, supplementary data and videos are available via FigShare at DOI 10.6084/m9.figshare.29503316.

Code: Used scripts are available in Github (https://github.com/AlexGalvez/discordances_in_phylogeny_and_morphology_within_loricat e_choanoflagellates.git).

Cell cultures and SEM stubs: A cell culture containing BEAP0094 (and mixed bacteria) is available at the Roscoff Culture Collection with identifier RCC11335, and also at the Culture Collection of Algae and Protozoa (CCAP) with identifier CCAP 2950/1. The corresponding SEM stub is deposited in the Marine Biological Reference Collections (CBMR) at the Institut de Ciències del Mar (ICM-CSIC, Barcelona, Spain) under the catalog/accession number ICMCBMR000696 (Guerrero et al., 2023). A cell culture containing BEAP0360 and mixed bacteria (also known as SIOpierAcanth1 and 10tr) is publicly available at the CCAP with identifier CCAP 2954/1. The corresponding SEM stub is deposited in the Marine Biological Reference Collections (CBMR) at the Institut de Ciències del Mar (ICM-CSIC, Barcelona, Spain) under the catalog/accession number ICMCBMR000697 (Guerrero et al., 2023) .

## CONFLICT OF INTEREST

The authors declare they have no conflict of interest relating to the content of this article.

## ACKNOWLEDGEMENTS

We thank Clara Cardelús, operating the Blanes Bay Microbial Observatory (BBMO), and Drs. Josep M. Gasol and Ramon Massana, for providing access to monthly water samples from Blanes Bay. We are grateful to Cédric Berney for discussions on the ribosomal sequences of the two choanoflagellate isolates, Martin Carr, Helge Thomsen and Barry Leadbeater for discussions on loricate taxonomy, and Mercedes Arrúe and Eduard Cortés for assistance with sample logistics.

## FUNDING

This project has received funding from the European Research Council (ERC) under the European Union’s Horizon 2020 research and innovation programme (grant agreement No. 949745), from the Departament de Recerca i Universitats de la Generalitat de Catalunya (exp. 2021 SGR 00751), from the grants QC2021-007134-P and PID2023-152955NA-I00 funded by MICIU/AEI/10.13039/501100011033 and by ERDF/EU. FUR is supported by the Howard Hughes Medical Institute Hanna H. Gray Fellows Program.

## Notes

https://figshare.com/articles/dataset/Data_for_b_Looks_can_be_deceiving_discordances_in_phylogeny_and_morphology_within_loricate_choanoflagellates_b_/29503316

https://github.com/AlexGalvez/discordances_in_phylogeny_and_morphology_within_loricate_choanoflagellates.git

### Competing Interest Statement

The authors have declared no competing interest.

