## Supplementary Figures for "Looks can be deceiving: discordances in phylogeny and morphology within loricate choanoflagellates"

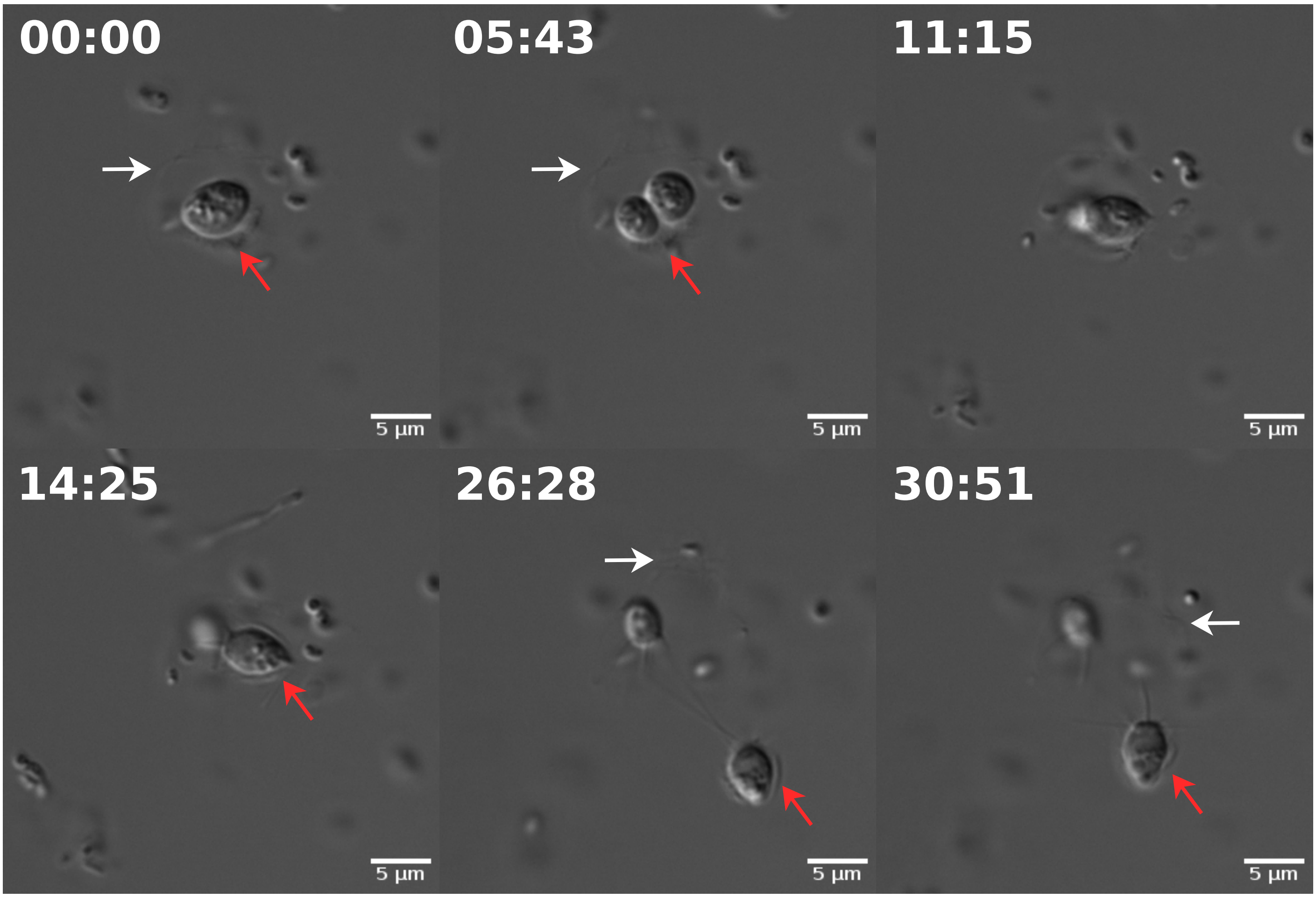


**Supplementary Figure 1. BEAP0094 portraying tectiform division. 00:00.** Loricate cell preparing for division, with costae already formed beneath. **05:43.** Cytokinesis. **11:15.** Daughter cell initiates inversion and retrieval of costae. **14:25.** Daughter cell enters the lorica previously constructed by the mother cell. **26:28.** Daughter cell exits the maternal lorica using a couple of threads. **30:51.** Daughter cell migrates using filopodia. White arrows indicate the mother lorica, while red arrows indicate the daughter lorica. Time-point format is mm:ss.

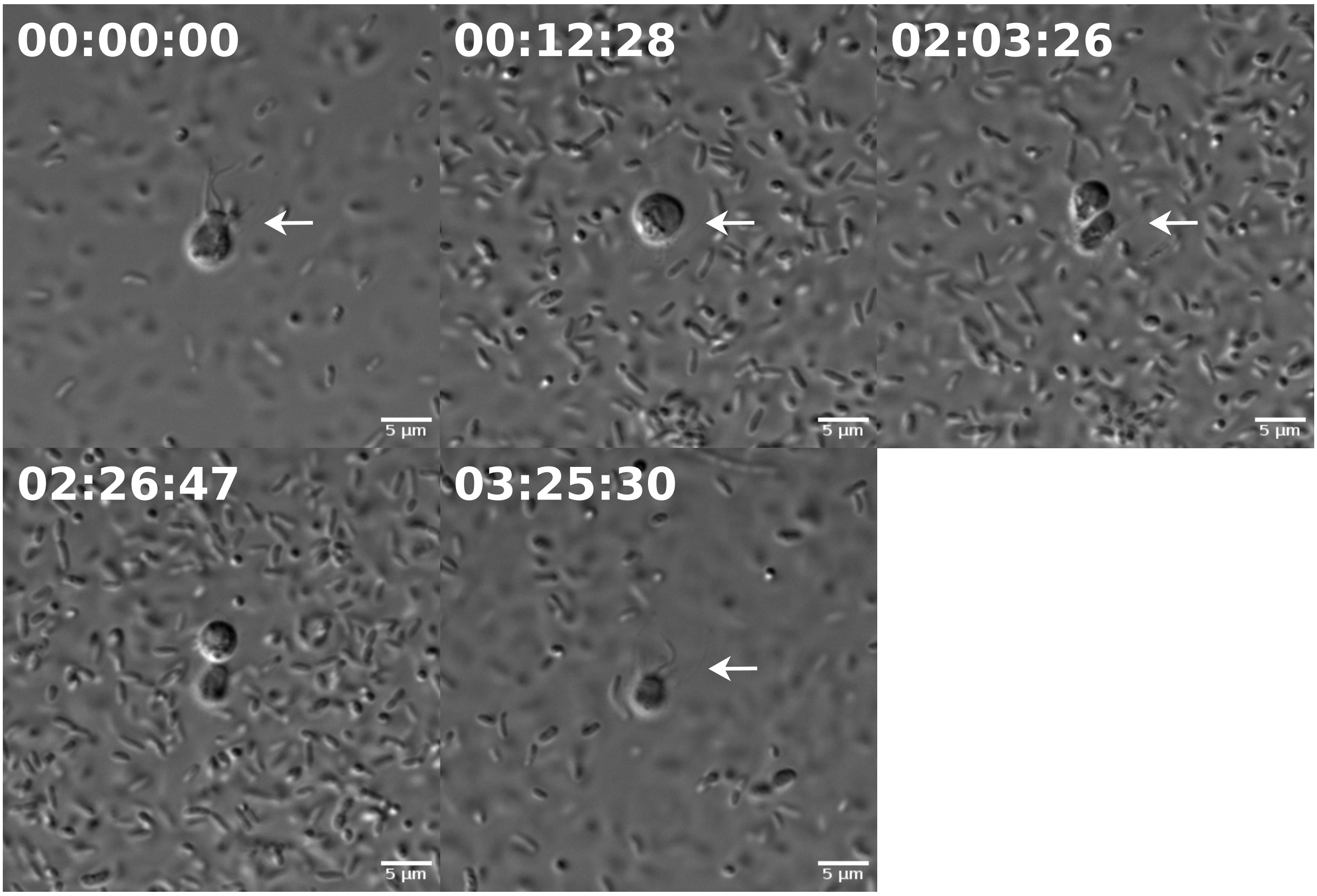


**Supplementary Figure 2. BEAP0360 portraying nudiform division. 00:00:00.** Loricate cell preparing for division. **00:12:28.** Loricate cell moves from the anterior to the posterior chamber. **02:03:26.** Diagonal cytokinesis **02:26:47.** Mother cell re-enters the anterior chamber. **03:25:30.** Daughter cell swims away before generating its own lorica. White arrows indicate the mother lorica. Time-point format is hh:mm:ss.
